# Rapid covalent-probe discovery by electrophile fragment screening

**DOI:** 10.1101/442806

**Authors:** Efrat Resnick, Anthony Bradley, Jinrui Gan, Alice Douangamath, Tobias Krojer, Ritika Sethi, Anthony Aimon, Gabriel Amitai, Dom Belini, James Bennett, Michael Fairhead, Oleg Fedorov, Paul P. Geurink, Jingxu Guo, Alexander Plotnikov, Nava Reznik, Gian Filippo Ruda, Laura Diaz Saez, Verena M. Straub, Tamas Szommer, Srikannathasan Velupillai, Daniel Zaidman, Alun R. Coker, Christopher G. Dowson, Haim Barr, Killian V.M. Huber, Paul E. Brennan, Huib Ovaa, Frank von Delft, Nir London

## Abstract

Covalent probes can display unmatched potency, selectivity and duration of action, however, their discovery is challenging. In principle, fragments that can irreversibly bind their target can overcome the low affinity that limits reversible fragment screening. Such electrophilic fragments were considered non-selective and were rarely screened. We hypothesized that mild electrophiles might overcome the selectivity challenge, and constructed a library of 993 mildly electrophilic fragments. We characterized this library by a new high-throughput thiol-reactivity assay and screened them against ten cysteine-containing proteins. Highly reactive and promiscuous fragments were rare and could be easily eliminated. By contrast, we found selective hits for most targets. Combination with high-throughput crystallography allowed rapid progression to potent and selective probes for two enzymes, the deubiquitinase OTUB2, and the pyrophosphatase NUDT7. No inhibitors were previously known for either. This study highlights the potential of electrophile fragment screening as a practical and efficient tool for covalent ligand discovery.

## Introduction

Targeted covalent inhibitors have many advantages as chemical probes and drug candidates ^1, 2^. These include: prolonged duration of action ^3^, improved potency and exquisite selectivity when targeting non-conserved protein nucleophiles ^4^. For these reasons and more, there has been increasing interest in covalently acting compounds, in both academia ^5^ and the pharmaceutical industry ^5–8^. This trend is underlined by the recent FDA approvals of the rationally-designed covalent drugs ibrutinib, afatinib, osimertinib and neratinib.

Discovery of new covalent inhibitors remains challenging, however. Historically, the most widespread approach for the design of such inhibitors relied on the incorporation of an electrophile into an already optimized reversible recognition element ^4, 9–11^, most notably in kinase inhibitors ^4, 12–16^. More recently, large-scale covalent virtual screens have also emerged as a method for the discovery of covalent binders ^17–22^. While successful, *in silico* docking still has its limitations: it is limited to targets for which a crystal structure (or a high-quality model) is available; and second, it cannot efficiently address protein flexibility. Empirical high throughput screening (HTS) for covalent binders is typically avoided ^23^ owing to concerns about promiscuous activity ^24–26^. A major risk in screening large covalent libraries is that hits will be dominated by overly reactive compounds rather than by specific recognition ^27^.

Fragment based screening, which focuses on very low molecular-weight compounds, is a successful hit discovery approach for reversible inhibitors ^28, 29^, that has led to several drugs and chemical probes ^29, 30^. Compared to traditional HTS, fragment-based screening offers better coverage of chemical space and higher probability of binding due to lower complexity ^31, 32^. The major limitation in fragment-based screening is the weak binding affinity of fragment hits, which not only necessitates very sensitive biophysical detection methods, coupled with elaborate validation cascades to eliminate attendant artefacts, but additionally makes progressing hits to potency difficult and expensive. In particular, it requires large compound series with typically ambiguous structure-activity relationships, because no method to date can reliably rationalize which are the dominant interactions of the original fragment. Screening *covalent* fragments addresses both problems: covalent binders are easy to detect by mass spectrometry; and because the dominant interaction is unambiguous, namely the covalent bond, designing follow-up series is simplified, and the primary hits are already potent.

A prominent covalent fragment screening approach is disulfide tethering ^33, 34^, which entails incubating a library of disulfide-containing fragments with the target. Disulfide exchange with the target cysteine selects for fragments that are reversibly stabilized in its vicinity. Disulfide tethering was successfully applied to a variety of targets containing both native and introduced cysteine residues ^35^. Recently it led to the discovery of a promising K-Ras^G12C^ inhibitor ^36^. Disulfides are not, however, suitable as cellular probes, and replacing them with a suitable electrophile is in general no less challenging than starting from a reversible ligand.

A potential solution is to directly screen mild electrophile fragments. Electrophile fragment screens were recently performed in small scale, with libraries of up-to ~100 compounds *in vitro* against a recombinant target ^37–41^ or in a cellular phenotypic context ^42–44^. Small scale screens were also performed with reversible covalent fragments ^45, 46^. We hypothesized that significantly increasing the library size and screening it against a diverse panel of targets will allow robust discovery of covalent ligands.

Here, we report a holistic covalent fragment screening approach. We have screened approximately 1000 electrophiles against ten different proteins. Combined with a newly reported high-throughput thiol-reactivity assay, our approach circumvents problems ascribed to irreversible binding: by robust evaluation of reactivity, and screening many proteins, we could detect and thus avoid promiscuous hitters. We could thus exploit the advantages of covalent screening, namely sensitive detection of binding at relatively low concentrations, yielding potent and selective primary hits in the majority (7 out of 10) of the cases. Moreover, we demonstrate that by combining the approach with high-throughput crystallography, quality leads can be rapidly developed, as shown for OTUB2 and NUDT7, two targets that previously lacked probes.

## Results

### Assembling an electrophile fragment library

We constructed our electrophile fragment library, by focusing on two mild electrophilic ‘warheads’: acrylamides and chloroacetamides. Acrylamides are represented in many rationally designed covalent drugs such as ibrutinib and osimertinib. Although chloroacetamides are more reactive ^47^, they still show selectivity in chemical-proteomic screens ^42^. Not many fragments containing these electrophiles were available for purchase from commercial vendors, likely due to the longstanding bias against covalent modifiers ^24, 25^. Nevertheless, we limited our screen to commercially available compounds, to enable the broadest future use of the library. Additional considerations were an overall low molecular weight, and enriching the library with related analogs to allow preliminary structure activity relationships to be deduced directly from a primary screen. The final library (Fig. 1) contains 993 compounds, comprising 76% chloroacetamides (n=752) and 24% acrylamides (n=241). 92% of the compounds have a molecular weight below 300 Da, including, in the case of chloroacetamides, the chlorine leaving group (36 Da). Thus, the molecular weight distribution of the reversible recognition elements is shifted to even lower masses (See Supp. Fig. 1 for molecular weight distribution in absence of the acrylamide/ choloroacetamide). 95% of the compounds have fewer than 20 heavy atoms, if the electrophilic moiety is included. The library also adheres to the so-called ‘rule of three’ ^48^, with almost all compounds containing fewer than three hydrogen-bond donors, acceptors, rotatable bonds and a cLogP <3 (Fig. 1).

**Figure 1.**
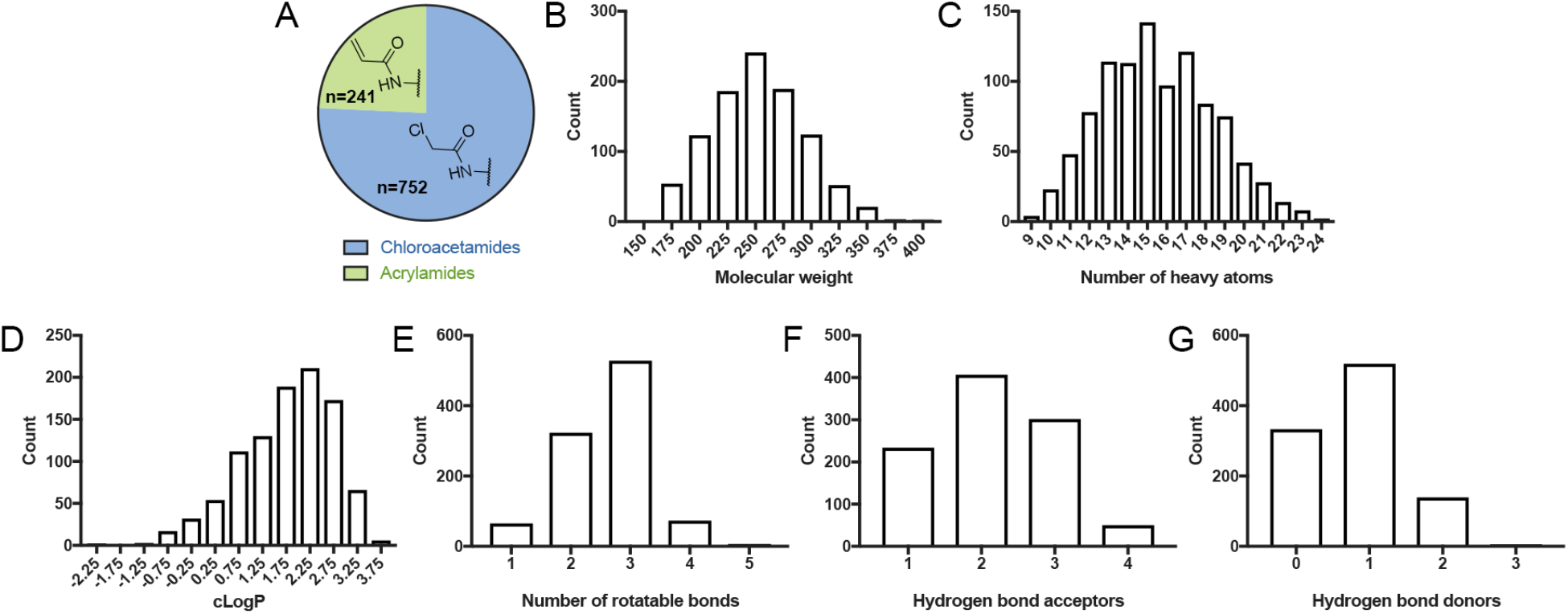
Electrophile fragment library adheres to the ‘rule-of-three’. Distribution of: **A.** Chloroacetamides and acrylamides in the library **B.** Molecular weights of the fragments, including the electrophile moiety **C.** Number of heavy atoms **D.** cLogP values **E.** Number of rotatable bonds **F.** Number of hydrogen-bond acceptors **G.** Number of hydrogen-bond donors. The library largely adheres to the “rule-of-three” for fragment libraries.

### Only a few fragments are highly reactive

To address a major concern in covalent-molecule screening, namely that high reactivity would lead to a high proportion of irrelevant hits, we developed a high-throughput thiol-reactivity assay in order to assess the reactivity of all fragments in our library. This entails incubating fragments with reduced DTNB (Ellman’s Reagent; 5,5-dithio-bis-2-nitrobenzoic acid) and following the absorbance of TNB^2-^ (at 412 nm wavelength) for up to seven hours. By fitting the data to a second order reaction rate equation we could extrapolate the kinetic constant for the alkylation (See example in Fig. 2; Supp. Dataset 1). The majority of the compounds showed an excellent fit to the kinetic model (63.5% had R ^2^>0.9; 71% had R ^2^>0.8; Supp. Fig. 22). The poorest-fitting data were obtained for the least reactive compounds (Supp. Fig. 22) for which the reaction rate was below the dynamic range of the assay.

Several results arise from the analysis of the kinetic data for the entire library. First, the most reactive fragments labeled only ~100 fold faster than the least reactive fragments. The majority of the library (78%) falls within only a 30-fold difference. This relatively narrow range of reactivity suggests that it is feasible to compare the labeling of fragment electrophiles in a screening scenario. Second, the chloroacetamides are clearly more reactive (show faster kinetics) than the acrylamides (*p*=4.6×10^−38^; two sided t-test), in accordance with previous anecdotes for a handful of fragments ^47^. Finally, for reference, we measured the reactivity of iodoacetamide, a generally used, non-selective thiol alkylator, and found it to be 16-fold more reactive than the average chloroacetamide, and 19-fold more reactive than the median choloroacetamide (Supp. Dataset 1).

**Figure 2.**
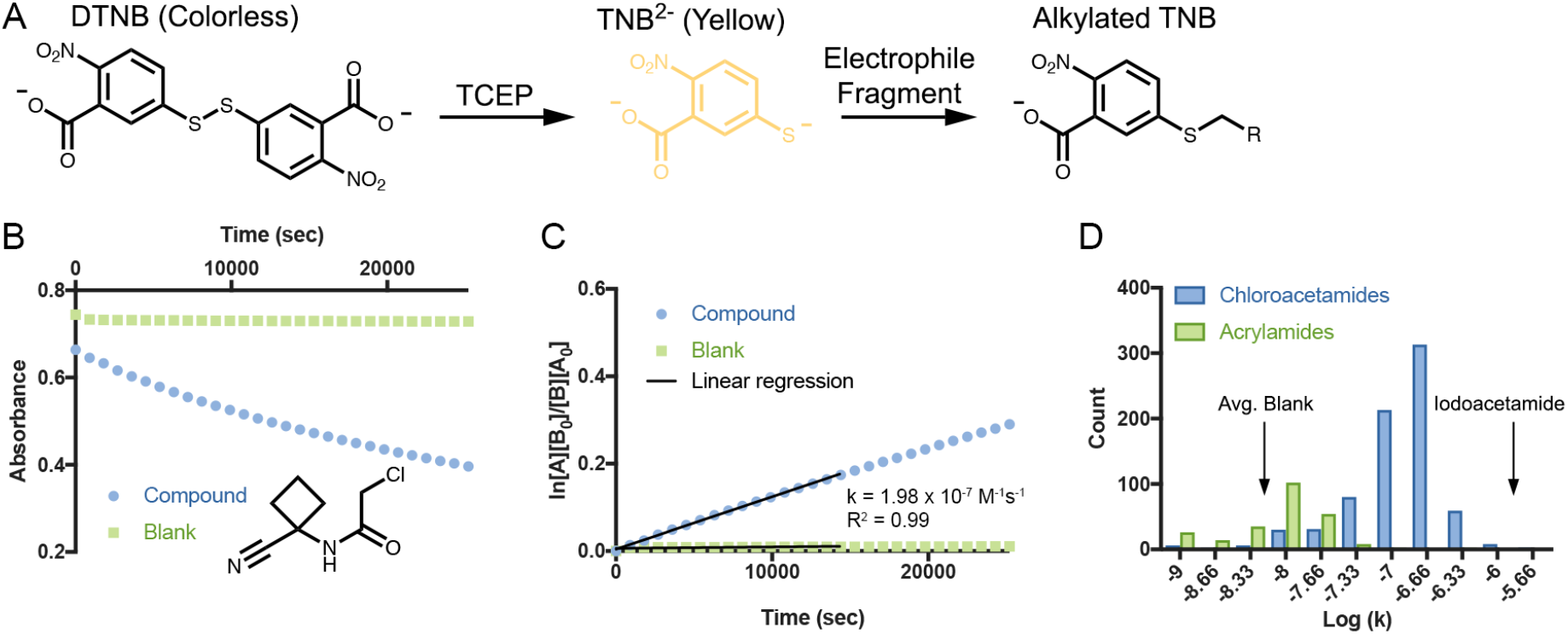
Narrow range of thiol-reactivity across electrophile fragments. **A.** Schematic description of the high-throughput thiol reactivity assay. In the presence of TCEP, DTNB is reduced to TNB^2-^ which has strong absorbance at 412 nm and is yellow under natural light. Alkylation of TNB^2-^ by an electrophile fragment reduces the observed absorbance. **B.** Example of the reactivity measurement for PCM-0102854. **C.** Example of second order kinetic rate calculation. The data is fitted to a second order reaction, [A] is the concentration of the electrophile and [B] is the concentration of TNB^2-^. The rate is determined by a linear regression of the data across four hours of measurement. **D.** Distribution of the rates of all electrophile fragments in the library, shows mild reactivity and narrow variability.

We therefore conclude, that while there is variability in the intrinsic thiol-reactivity of the fragments, it is sufficiently small to allow the identification of quality hits in a screening campaign. Moreover, these data indicate that the two selected electrophiles are indeed sufficiently mild to ensure the main driver of protein labeling is recognition rather than reactivity.

### Intact protein mass spectrometry identifies hits against a diverse panel of proteins

We screened our library against a diverse panel of ten cysteine-containing proteins (Table 1). The targets were selected based on their therapeutic potential, and most of them lacked any validated covalent inhibitor or probe, or indeed any known chemical probe at all. As a control, we screened Bovine Serum Albumin (BSA). Four of the proteins in the panel contain a solvent exposed catalytic cysteine, while the other six do not.

Each protein was incubated with the electrophilic library in pools of five compounds per well, each at 200 μM, for 24 hours at 4 °C to allow screening of proteins that are not stable at higher temperatures for long time periods. Following incubation, we used intact protein liquid chromatography/mass-spectrometry (LC/MS) to identify and quantify labeling by the fragments (Fig. 3A). Overall, while the hit rate varied greatly between different proteins (Fig. 3B), we were able to find hits for almost all of the screened proteins, except for PCAF, QSOX1 and the negative control BSA (See Supp. Dataset 2 for the labeling quantification; Supp. Fig. 3 for structures of selective hits).

**Table 1.** Panel of protein targets for screening.

| Protein<sup>a</sup> | MW<sup>b</sup> | Hits<sup>c</sup> | Selective Hits<sup>d</sup> | Catalytic Cys<sup>e</sup> | Cys residues<sup>f</sup> | Possible therapeutic indications<sup>g</sup> |
| --- | --- | --- | --- | --- | --- | --- |
| QSOX1 | 58,084 | 0/993 | 0 | No<sup>h</sup> | 12 | Cancer<sup>79,80</sup> |
| PCAF | 19,452 | 0/993 | 0 | No | 3 | HIV<sup>81</sup><br>Cancer<sup>82,83</sup> |
| PBP<sup>R504C</sup> | 57,903 | 2/983 | 2 | No | 1 | Antibiotic resistance<sup>84</sup> |
| K-Ras<sup>G12C</sup> | 19,245 | 10/968 | 7 | No | 3 | Cancer<sup>36,85</sup> |
| USP8 | 44,429 | 20/923 | 7 | Yes | 12 | Cancer<sup>86</sup><br>Cushing's disease<sup>87</sup> |
| NNMT | 31,248 | 30/299 | 22 | No | 8 | Cancer<sup>88</sup><br>Diet-induced obesity<sup>89</sup> |
| OTUB2 | 27,312 | 47/938 | 42 | Yes | 4 | Viral infection<sup>53</sup><br>Diabetes<sup>54</sup><br>ALS<sup>55</sup> |
| NUDT7 | 26,672 | 36/973 | 24 | Yes | 4 | Diabetes<sup>61</sup> |
| NV3CP | 19,284 | 10/824 | 9 | Yes | 5 | Viral infection<sup>72</sup> |
| BSA | 66,464 | 0/981 | 0 | No | 23 | Negative control |

<sup>a</sup> Acronyms are as follows: Quiescin sulphhydryl oxidase 1 (QSOX1); Penicillin-binding protein 3 (PBP); Ubiquitin-specific peptidase 8 (USP8); Nicotinamide N-methyltransferase (NNMT); Norovirus 3C protease (NV3CP); Bovine serum albumin (BSA).
<sup>b</sup> Molecular Weight of construct in Da
<sup>c</sup> Number of fragments with >50% labeling / compounds for which labeling could be evaluated
<sup>d</sup> Number of hits that did not label any other protein to >50%
<sup>e</sup> Indicates whether a catalytic cysteine is present in the protein
<sup>f</sup> Number of cysteine residues in protein
<sup>g</sup> Indicates possible therapeutic rationale to inhibit this target.
<sup>h</sup> QSOX1 contains a catalytic disulfide.

**Figure 3.**
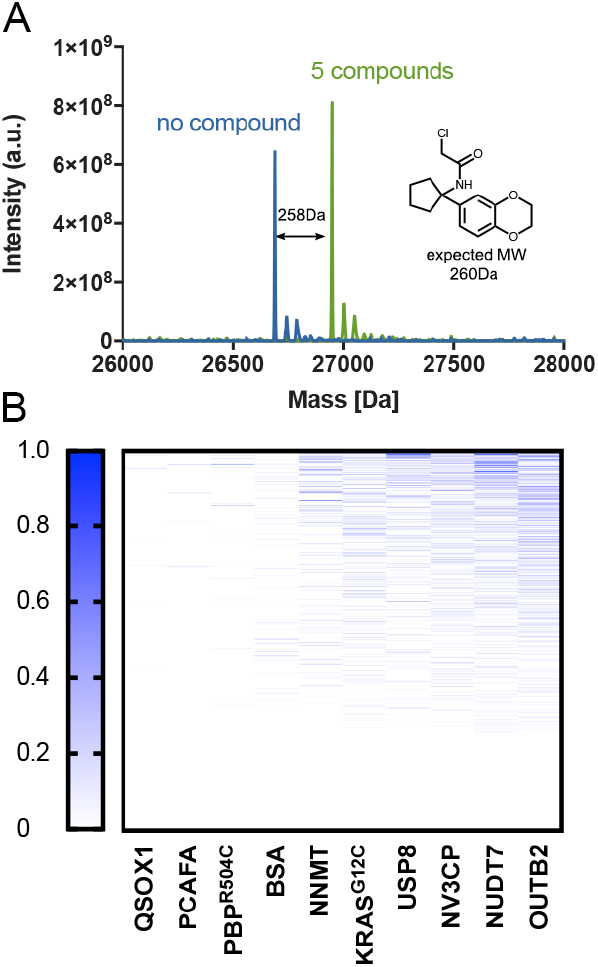
Intact protein LC/MS screen identifies hits for most targets. **A.** An example of the LC/MS deconvoluted spectrum for NUDT7 with no compound (blue) and after 24 hours incubation with five compounds (green), the shift in the mass of the protein corresponds to 100% labeling of PCM-0102951. **B.** Summary of the quantified labeling of ten proteins by the electrophile library. Blue represents 100% binding and white no labeling or data not available (see labeling assignment in methods).

### Promiscuity does not correlate with reactivity

We define promiscuous compounds to be those that label two or more proteins by more than 50%, or three proteins by more than 30%. Despite this stringent definition, only 27 of the electrophilic fragments are promiscuous (Supp. Fig. 4; Supp. Dataset 2). Under an even stricter definition, of more than 30% labeling of any two proteins, only an additional 36 compounds become promiscuous.

Unexpectedly, promiscuous labeling does not correlate well with thiol-reactivity (Supp. Fig. 5; R ^2^=0.09). For instance, compound PCM-0102496 (Supp. Fig. 4) labeled NNMT, USP8 and NUDT7 by more than 50%, although its alkylation rate is in the lowest quartile of reactivity (1.09×10^−8^ M^−1^s^−1^). On the other hand, some of the most reactive compounds such as PCM-0102140, PCM-0102859, PCM-0102150 (Supp. Fig. 55), do not label any protein at all.

We evaluated the possibility that promiscuous compounds label amino acids other than cysteines. We incubated five promiscuous compounds with NUDT7 and USP8 followed by trypsin digestion and LC/MS/MS analysis to identify modification sites. Despite rare lysine and histidine modifications, the compounds preferably reacted with cysteines (Supp. Dataset 4) largely ruling out this hypothesis.

Degradation of compounds between protein screening and reactivity measurement can also explain the discrepancy between promiscuity and reactivity. To control for this, we re-sourced 14 compounds - ten of the most promiscuous compounds and four random compounds that did not label any protein. We evaluated the reactivity of these fresh compounds again and with the exception of one compound (PCM-0102982) the rates agreed well with the previous measurements (Supp. Table 1), suggesting degradation is not the source of the discrepancy.

Many of the promiscuous binders contain similar chemical motifs. For instance, we identified a large family of aminothizole chloroacetamides (Supp. Fig. 55; Supp. Fig. 6) that are frequent hitters in our screens. These are not significantly more reactive than other chloroacetamides in our thiol-reactivity assays (*p*=0.183 in a one sided t-test).

### Fragment growing identifies novel OTUB2 inhibitors

OTUB2 is a deubiquitinase (DUB) from the ovarian tumor domain (OTU) DUB superfamily ^49^. OTUB2, initially identified in HeLa cells ^50^, preferentially cleaves Lys63-linked polyUb chains and can also cleave Lys11- and Lys48-linked chains ^51^. OTUB2 is important to the choice between the homologous recombination and the nonhomologous end joining DNA repair pathways ^52^. OTUB2 has also been found to function as negative regulator of virus-triggered type I IFN induction ^53^, and was linked to inhibition of NF-κB signaling and regulation of beta cell survival in human pancreatic islets ^54^. Finally, OTUB2 has been identified as potential biomarker for sporadic amyotrophic lateral sclerosis (ALS) ^55^. As such, OTUB2 plays an important role in several biological pathways and the development of OTUB2-specific inhibitors can have therapeutic potential.

The primary screening against OTUB2 produced 47 fragments with >50% labeling, of which 39 were non-promiscuous and 37 strictly non-promiscuous (Supp. Fig. 7). We evaluated protein labeling of 26 of these compounds, at 200 and 100 μM (24 hours; 4 °C; Supp. Table 2), and nine compounds showed >50% labeling even at the lower concentration. In order to prioritize fragments for optimization we turned to high-throughput crystallography.

We were able to determine co-crystal structures of 15 OTUB2/fragment complexes by streamlined parallel co-crystallization involving 24 hours of pre-incubating protein with fragment and seeding with apo-protein crystals. In 11 of these complexes the fragments formed a covalent bond with the catalytic cysteine 51 in the enzyme active site (Fig. 4A; Supp. Table 3). The carbonyls of all chloroacetamides occupied the oxyanion hole formed by the amide backbones of D48, G49, N50, and C51. To progress selected fragments, analysis of labeling results and crystal structures led us to focus on a series of fragments sharing a common chloroacethydrazide motif (Fig. 4B), two of which were seen in crystal structures. In both cases the shared moiety participates in an extensive hydrogen bonding network with the protein active site (Supp. Fig. 8). The side chain of E174 switched its rotamer (in comparison of all other apo and fragment bound co-crystal structures) in order to mediate one such hydrogen-bond to the hydrazide motif (Supp. Fig. 9). In both structures, the hydrophobic moiety connected to the hydrazide pointed towards the solvent, making few obviously productive contacts with the protein. We concluded we might be able to optimize compound binding by changing this moiety.

**Figure 4.**
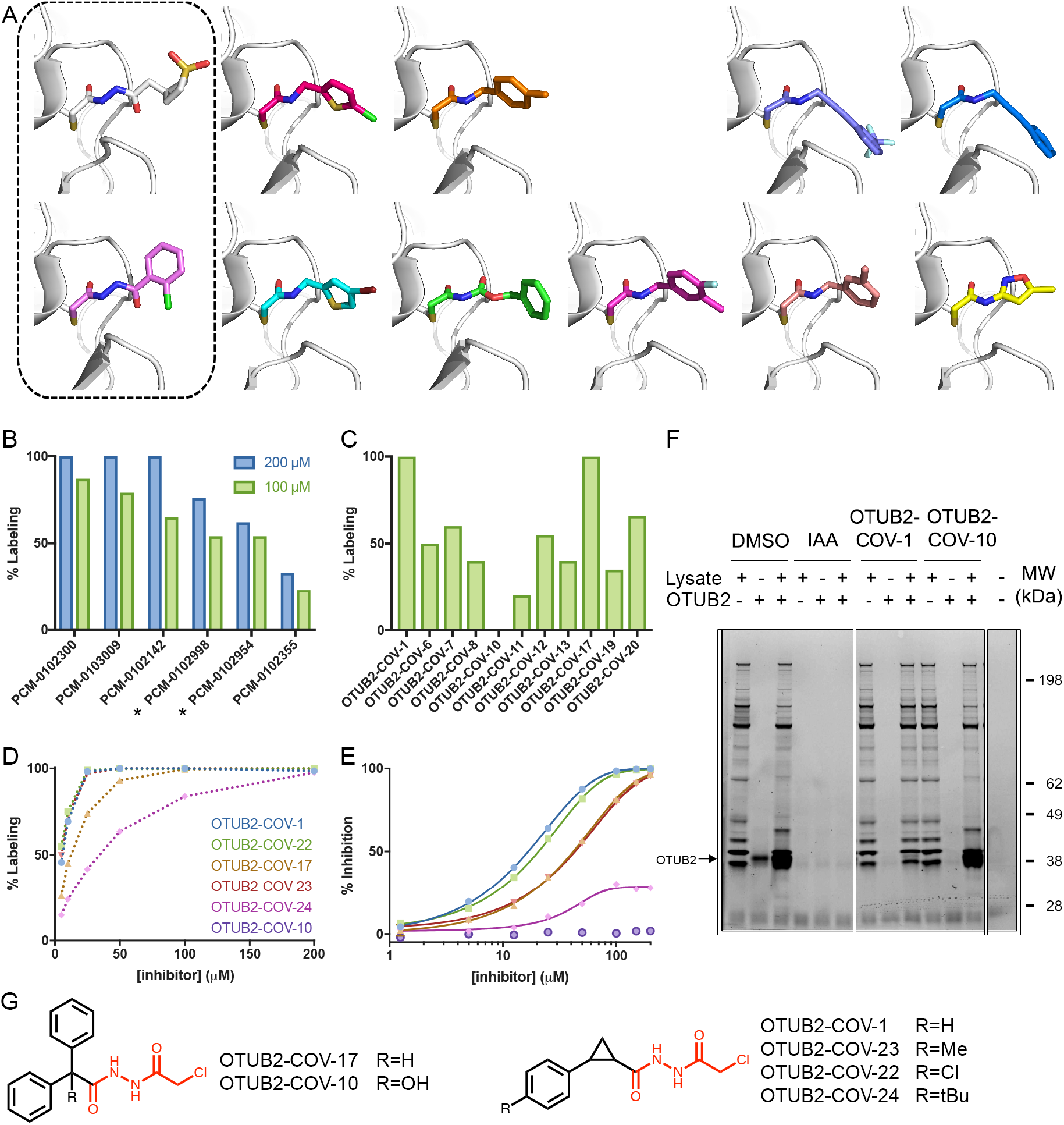
Discovery of a selective OTUB2 inhibitor by fragment growing. **A.** Co-crystal structures of OTUB2 in complex with (from top left) PCM-0102998, PCM-0102973, PCM-0102660, PCM-0103011, PCM-0103007, PCM-0102954, PCM-0103050, PCM-0102153, PCM-0102305, PCM-0102821, PCM-0102500 (See Supp. Fig. 7). Structures with compounds containing the chloroacethydrazide motif are boxed. **B.** % covalent labeling of OTUB2 with compounds containing the identified motif, PCM-0102300, PCM-0103009, PCM-0102142, PCM-0102998, PCM-0102954, PCM-0102355 (Supp Fig. 7) at 200 μM (blue) and 100 μM (green). Compounds boxed in (A) are marked with asterisks **C.** % covalent labeling of OTUB2 with selected next generation compounds at 100 μM (See Supp Fig. 10 for all analogs, and Supp. Table 4 for % labeling). **D.** Dose response measurement of % labeling by next generation OTUB2 binders. All labeling in (B-D) are measured after 24 hours incubation in 4 °C. **E.** Inhibition of OTUB2 in an enzymatic assay (2.5 hours pre-incubation in the presence of 2 mM free cysteine). **F.** Chemical proteomics selectivity assessment using a fluorescent activity-based DUB probe ^56^. Lanes 1-3: DMSO negative controls showing probe labeling of DUBs in lysate, of purified OTUB2 and lysate spiked with OTUB2 (0.05 μg). Lanes 3-6: Iodoacetamide (10mM) as positive control eliminates all probe labeling. Lanes 7-9: OTUB2-COV-1 specifically compete with probe only for OTUB2. Lanes 10-12: Negative control compound OTUB2-COV-10 can only compete with probe for purified OTUB2 but not for OTUB2 in lysate. Note, it does have other DUB off-targets (compare lanes 10 and 12) **G.** Chemical structures of selected next generation OTUB2 binders. Chloroacethydrazide motif is highlighted in red.

We purchased 21 analogs, all containing the chloroacethydrazide motif (Supp. Fig. 10). When incubated with OTUB2 at 100 μM (24 hours; 4 °C), six showed >50% labeling, representing the reversible recognition stemming from the shared motif (Fig. 4C; Supp. Table 4). Two particularly promising analogs, OTUB2-COV-1 and OTUB2-COV-17, showed 100% labeling (Fig. 4C). We sourced additional analogs of OTUB2-COV-1 with various *para* substitutions of the phenyl ring, and assessed their labeling efficiency at various concentrations. Compounds OTUB2-COV-1, OTUB2-COV-22 and OTUB2-COV-23 showed 46% and 55% and 49% labeling respectively at 5 μM (24 hours; 4 °C; Fig. 4D).

We next assessed their inhibition of the enzymatic activity of OTUB2 (Fig. 4E). The best inhibitor, OTUB2-COV-1 showed an IC_50_ of 31.5 μM at a relatively short incubation time of 30 minutes and in the presence of 2 mM cysteine. The IC_50_ improved to 15.4 μM with a longer incubation of 2.5 hours, supporting the irreversible binding mechanism. There was a very good correlation between the labeling efficiency of these compounds and their inhibitory effect. We attempted to determine a co-crystal structure of OTUB2 labeled with the lead compound OTUB2-COV-1. While the chloroacethydrazide motif adapted the same conformation as the original library hits, we could not detect density for the cyclopropyl-phenyl moiety (Supp. Fig. 11).

The improvement of the analogs appears to stem from better recognition rather than reactivity: the primary chloroacethydrazide hits gave little labeling of other proteins in the panel (Supp. Dataset 2), and their mild reactivity in the thiol-reactivity assay (Average Rate k=3.9×10^−8^ M^−1^s^−1^), was comparable to that of some of the new analogs in the same assay (OTUB2-COV-23 k=2.74×10^−8^ M^−1^s^−1^; OTUB2-COV-22 k=3.69×10^−8^ M^−1^s^−1^; Supp. Dataset 1).

To further show the compounds’ selectivity and to validate labeling in complex mixtures we assessed their promiscuity against all DUBs in HEK293 lysates, by chemical proteomics (Fig. 4F). We used our previously developed fluorescent activity-based DUB probe ^56^ to label all DUBs in lysates pre-incubated (3 hours, 37 °C) with either DMSO (control) or compound (50μM). OTUB2 is endogenously expressed at very low levels and so is not visible compared to other highly expressed DUBs. We thus performed the same experiment with a recombinantly expressed OTUB2, and with lysates spiked with 0.1 or 0.05 μg OTUB2.

Strikingly, even at such high compound concentration, there were no detectable differences between the compound-treated lysate and DMSO control (Fig. 4F, compare lanes 1 and 7), indicating exquisite selectivity across all DUBs detected by the probe. When incubated with OTUB2 alone, the fragments outcompeted the DUB probe, completely blocking any labeling at 0.05 μg (Fig. 4F, lane 8) and significantly diminished labeling at 0.1 μg (Supp. Fig. 12C). This effect is much more pronounced in spiked lysates. A very pronounced band appears for OTUB2 in the DMSO control as well as the inactive compound control OTUB2-COV-10 (Fig. 4F, lanes 3 and 12; likely due to merging with bands of close molecular-weight DUBs). However, for the most potent compound OTUB2-COV-1, this band completely disappears (Fig. 4F, lane 9). For the other analogs, while all diminish the probe labeling against recombinant OTUB2, none are able to compete with it as well in the spiked lysate (Supp. Fig. 12A).

### Fragment merging leads to potent NUDT7 inhibitors

NUDT7 is a peroxisomal CoA pyrophosphohydrolase and belongs to a protein family characterised by a 23-amino acid motif referred to as the ‘NUDIX box’. These proteins have been reported to hydrolyse a diverse range of substrates including (d)NTPs, nucleotide sugars, diadenosine polyphosphates as well as capped RNA ^57^. The NUDT7 gene contains a CoA-binding motif and a C-terminal peroxisomal targeting signal (PTS) ^58, 59^. Expression of NUDT7 is highest in liver, with NUDT19 likely acting as the complementary CoA and CoA ester hydrolase in kidney ^60^. Leptin double knockout mice, which display alterations in CoA homeostasis and exhibit a diabetic phenotype, have been reported to express reduced levels of NUDT7 with a concomitant increase in pantothenate kinase activity ^61^. To the best of our knowledge, no small molecule inhibitors or probes have been reported for NUDT7 so far.

The primary screening against NUDT7 produced 36 fragments with >50% labeling, of which 24 were non-promiscuous, and 20 strictly non-promiscuous (Supp. Fig. 13). A series of similar fragments sharing a common 2-phenylpyrrolidine motif stood out with four compounds labeling 100% (Fig. 5). We validated compound binding via differential scanning fluorimetry (DSF) in which 26 of the 30 compounds that showed covalent labeling also stabilized NUDT7 by 4.5 - 13.4 °C (Supp. Fig. 14). Specifically, compounds PCM-0102298, PCM-0102938, PCM-0102558, PCM-0102951, PCM-0102716 and PCM-0102512 stabilized NUDT7 by 4.5 - 8 °C (Fig. 5A; Supp. Fig. 13).

In order to optimize this series, we determined the co-crystal structures of compounds PCM-0102951, PCM-0102558 and PCM-0102716 bound to NUDT7 (Supp. Table 5) using a similar co-crystallization protocol as for OTUB2. The structures show that all compounds form a covalent bond with the catalytic cysteine 73. Surprisingly, however, despite their chemical similarity, the three compounds adopt different binding poses (Fig. 5D). Evaluation by dose-response labeling of the six compounds sharing the 2-phenylpyrrolidine motif showed that after incubation (4 °C, 24 hours), all six compounds label 100% up to 25 μM. At 5 μM concentration, the compounds label less (60-80%) except for PCM-0102558, which still shows 100% labeling (Fig. 5B).

**Figure 5.**
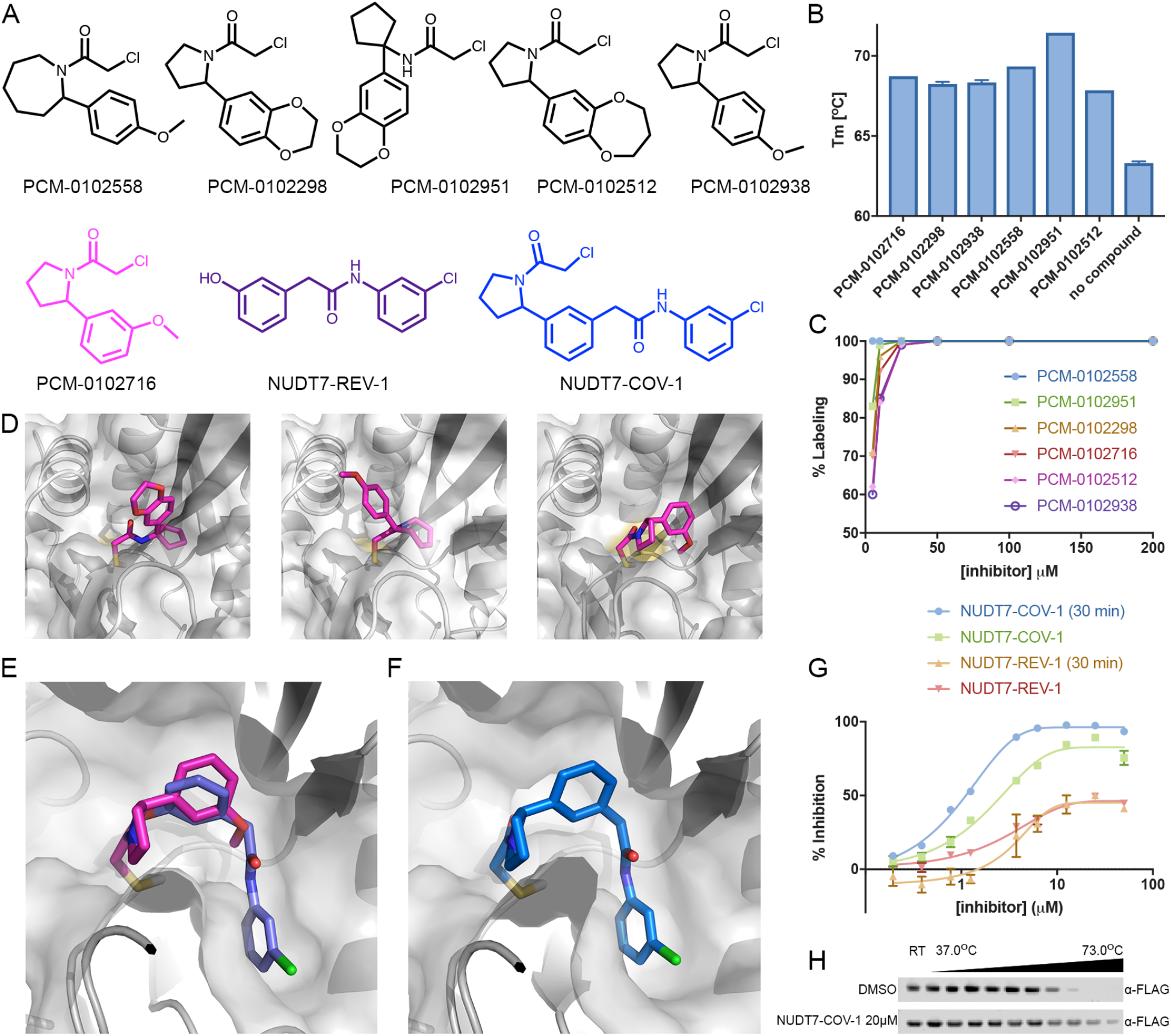
Discovery of a potent NUDT7 inhibitor by fragment merging. **A.** Chemical structures of similar hit compounds that labeled NUDT7 68% (PCM-0102716), 88% (PCM-0102512) and 100% (PCM-0102558, PCM-0102298, PCM-0102951, PCM-0102938) in the primary screen. Compound NUDT7-REV-1 is a non-covalent fragment (purple) that was identified as a NUDT7 binder in a crystallography soaking screen (See panel E). NUDT7-COV-1 (blue) is a merged compound based on PCM-102716 (magenta) and NUDT7-REV-1. **B.** The six hits identified in the primary screen stabilize NUDT7 by 4.5-8.1 °C in a T_m_ shift assay. **C.** Labeling percentage of compounds PCM-0102558, PCM-0102951, PCM-0102298, PCM-0102716, PCM-0102512 and PCM-0102938 at 5-200 μM. **D.** Co-crystal structures of NUDT7 with compounds PCM-0102951, PCM-0102558, PCM-0102716. **E.** Overlay of the crystal structures of NUDT7 with compound PCM-0102716 (pink) and with the non-covalent fragment NUDT7-REV-1 (purple). **F.** Co-crystal structure of NUDT7 with the merged compound NUDT7-COV-1 adopts the exact same binding mode as the two separate fragments. **G.** Enzymatic inhibition of NUDT7 by NUDT7-COV-1 and NUDT7-REV-1. Data shown includes results with and without protein incubation in the presence of the compounds. **H**. Intracellular target engagement is demonstrated by thermal stabilization of FLAG-NUDT7 by NUDT7-COV-1 in intact HEK293 cells. After transfection cells were treated with 20 μM NUDT7-COV-1 or DMSO for 30 minutes before heating to the indicated temperatures.

We had previously completed a crystallographic fragment screen with non-covalent fragments at the XChem facility at Diamond Light Source and identified 18 fragments bound to the putative substrate binding region of NUDT7 (10.5281/zenodo.1244111). Based on one of the initial hits, we synthesized a series of diphenyl-acetamide analogues and soaked them into NUDT7 crystals. This yielded a structure in complex with compound NUDT7-REV-1 (Fig. 5A), and a comparison of this structure with the covalent NUDT7/PCM-0102716 structure revealed an almost perfect overlap of one of the phenyl rings (Fig. 5E), suggesting a clear strategy for fragment merging.

The merged compound, NUDT7-COV-1 (See chemical synthesis in Supp. File 1), combines the key features of both fragments and has much improved properties. In the co-crystal structure, the merged compound adopts exactly the predicted pose (root mean square deviation of 0.4 Å over the shared atoms), and at 5 μM, NUDT7-COV-1 labels NUDT7 to 100% in 15 minutes (2 μM protein). This improvement is likely due to improved recognition since NUDT7-COV-1 is less than three-fold more reactive than its parent compound (k=4.22×10^−7^ M^−1^s^−1^ *vs.* k=1.63×10^−7^ M^−1^s^−1^ for PCM-0102716). We evaluated the selectivity of NUDT7-COV-1 in the same chemical proteomics experiment used for OTUB2 and observed no DUB off-target labeling, suggesting the compound does not display random off-target activity (Supp. Fig. 15).

In a NUDT7 enzymatic activity assay, without pre-incubation, the merged compound had an IC_50_ value of 2.7 μM. After 30 minutes pre-incubation with the compound the IC_50_ value improves to 1.1 μM. It is interesting to note that of the non-covalent hits from crystallographic fragment screening, none showed detectable activity in the enzymatic assay, including the parent non-covalent fragment we used for merging (Fig. 5G; Supp. Fig. 16; Supp. Fig. 17). Lastly, we evaluated NUDT7 cellular target engagement by NUDT7-COV-1 via a cellular thermal shift assay (CETSA) in intact HEK293 cells. Indeed, NUDT7-COV-1 showed significant stabilization of FLAG-tagged NUDT7 compared to DMSO, confirming the compound binds the target in living cells (Fig. 5H).

### Electrophile fragments can be suitable for cellular screens

Many previous studies established a correlation between intrinsic reactivity and cellular toxicity for a range of electrophiles ^62^. To see if such a correlation exists for our electrophilic fragments, we performed a cellular viability assay for each of our compounds with three different model cell lines, HEK293, HB2 and CCD841 (Supp. Dataset 3).

At a concentration of 10 μM and 48 hours incubation, 47% 58% and 60% of the compounds had negligible effect on viability of HEK293, HB2 and CCD841 cells respectively (>75% viable; Supp. Fig. 18A-C). There was good correlation between the toxicity of the compounds across the three cell lines (Supp. Fig. 18D-F). We see a switch like toxicity effect in which compounds with reaction rates of k=1×10^−7^ M^−1^s^−1^ or less hardly affect viability - only 30%, 19% and 18% of these compounds affect viability by more than 25% of HEK293, HB2 and CCD841 cells respectively. Whereas compounds with rates higher than k=1×10^−7^ M^−1^s^−1^ show sharp decline in viability as a function of reactivity (Supp. Fig. 18A-C). 89%, 80% and 75% of these compounds reduce cellular viability by more than 25%. These results indicate that this library can be suitable for cellular phenotypic screening in addition to *in vitro* screening against purified proteins.

## Discussion

Discovery of selective covalent acting compounds is challenging. We approach this problem by significantly increasing the chemical space of recognition elements, through use of mild electrophiles, with careful accounting for reactivity and promiscuity. We describe the screening of 993 commercially available electrophile fragments against ten different proteins. Previous electrophile screening campaigns were limited in scope: Nonoo, *et al.*^38^ assayed only ten acrylamides against three proteins; Jost *et al.*^40^ tested six diverse electrophiles against 11 proteins; Kathman *et al.*^37^ screened 100 methyl acrylates against four proteins; and most recently Craven *et al.*^39^ screened 138 electrophiles against the kinase CDK2. While these studies were restricted to well-studied targets with known inhibitors, they pioneered electrophilic fragment screening and served as a proof-of-concept that the method is viable. By significantly expanding the library and screening a diverse array of targets, we demonstrate the broad applicability of this approach by producing valuable hits for ‘orphan’ targets.

We observe, moreover, that the combination of screening carbon electrophiles alongside exploiting high-throughput crystallography, results in an effective method for progressing hits to leads and designing probes: the covalent bond in the hit provides a clear, dominant chemical rule for designing follow-ups, and provides sufficient potency such that only few analogs are necessary to pass the threshold for a potent and selective probe. Thus, fewer than 50 compounds were purchased or synthesized in achieving selective chemical probes against two targets.

Another key point of screening carbon electrophiles, especially compared to the well-established disulfide tethering approach, is that it directly optimizes both binding of a recognition element as well as the electrophile orientation towards the target cysteine. Disulfide hits from a tethering screen typically cannot be used as cellular probes due to the reducing environment, and the transition from an active disulfide fragment to a similarly active carbon electrophile can be demanding and require the synthesis of many test compounds. For example, Ostrem *et al.* ^36^ identified a tethering hit against K-Ras^G12C^, but had to synthesize nearly 100 carbon electrophiles to reach suitable labeling efficacy. Instead, our screen immediately identified compound PCM-0102818 (Supp. Fig. 3) which is highly similar to the previously optimized compounds, and labeled 63% of K-Ras^G12C^, highlighting the efficiency of directly screening carbon electrophiles. The expanded library size does however appear crucial to providing sufficient coverage of chemical space: our previous screen of K-Ras^G12C^ with a much smaller number of electrophiles ^18^, only 62 acrylamide fragments, failed to yield plausible hits.

A major result of this study is the relatively narrow range of reactivity displayed by the wide majority of the electrophilic fragments, an observation that is robust due to the large number of compounds we could screen with the new high-throughput reactivity assay. Previous studies that characterized the thiol-reactivity of various electrophiles often considered as warheads for chemical probes ^37, 47^, were limited in the number of evaluated compounds, likely due to the low-throughput assays used to determine thiol-reactivity. For example, when examining acrylamides, Kathman *et al.*^37^ determined the pseudo-first order reaction rate for only three model compounds, and because one showed a significantly higher reaction rate, suggested that acrylamides as a class of electrophiles have too variable reactivity for screening. Instead, we could assess the reactivity of close to 250 acrylamides. We too observed outliers with high reactivity, but the vast majority displayed low reactivity and narrow variability. Indeed, under the current conditions the rates of 90 acrylamides fall below the dynamic range of our assay, showing similar rates to the background blank reaction. Acrylamide hits were also rarer in the ten protein screens, overall suggesting that as a class of electrophiles, they are suitable for screening.

A related observation is that promiscuity is not in fact a function of reactivity, historically the main reason covalent compounds were assumed to be problematic. While promiscuous compounds were observed (Supp. Dataset 2) and will be removed from future screens, the most promiscuous compounds did not necessarily have high intrinsic reactivity. For instance, whereas PCM-0102957 (Supp. Fig. 4), one of the most promiscuous in the library, has an intrinsic reactivity similar to iodoacetmide (1.9×10^−6^ M^−1^s^−1^), another one, PCM-0102496 (Supp. Fig. 4), displays very low intrinsic thiol reactivity. A previous study that tried to correlate intrinsic reactivity of electrophiles to *in vitro* covalent binding found similar discrepancies ^63^. We discounted two possible explanations for this observation: that these unexpected compounds do not label cysteine, but other amino acids like lysine (Supp. Dataset 4); and that the low reactivities were an artefact of compounds degradation (Supp. Table 3). Other explanations such as photoreactivity, redox cycling and other confounding mechanisms might still be at play. This remains an area of active research, and future screens might shed more light on this phenomenon.

Nevertheless, the data allow us to recommend a threshold reactivity of k=1×10^−7^ M^−1^s^−1^, below which electrophiles are likely to be useful in screens, providing hits suitable for further optimization and progression to cell active probes. This is based on the loose correlation between reactivity and promiscuity (Supp. Fig. 5) and the correlation between reactivity and cellular toxicity (Supp. Fig. 18A-C). An exception to this rule of thumb would be structural motifs we identify as promiscuous such as the aminothiazole series (Supp. Fig. 6), which showed up as frequent hitters, even in non-covalent screening ^64^, and are considered PAINS compounds ^24^.

The hit rates obtained with this library are on-par or slightly higher than observed in screens with non-covalent fragments ^65, 66^: 2-4% for NNMT, OTUB2 and NUDT7; and 0.2-0.9% for other proteins. These would be attenuated by screening at different concentrations: here, all primary screening was performed at 200 μM, but based on the results, we can now recommend a concentration of 100 μM when targeting catalytic cysteines, while staying at 200 μM for less nucleophilic target cysteines.

A major application of our screening approach is the ability to evaluate potential ligandability of target cysteine residues. It is clear by looking at the overall labeling statistics (Fig. 3B) that different target cysteines show different potential for electrophilic labeling. Proteomic approaches for the identification of functional or reactive cysteines ^42, 44, 67–69^ can identify potential target cysteines in a much larger scale than ever before. However, the throughput of the proteomics pipeline does not allow to assess the covalent ligandability. As an example, in a recent proteomic screen of 50 electrophilic fragments, Backus *et al.*^42^ identified several cysteine residues available for labeling in NNMT, but did not deem it to be a probe target since only a single fragment was able to significantly label it. In contrast, in our library we found 30 compounds that labeled NNMT by more than 50%, and in an enzymatic assay with a much shorter incubation time (2 hours), ten of these were able to inhibit it (>20% inhibition; Supp. Dataset 5). Indeed, recently a report was published in which a selective probe could be developed against NNMT in lysates but not in whole cells ^70^.

Overall, we conclude our approach can be widely adopted: the screening requires no specialized equipment or algorithms, the compounds of the library are commercially available from a single vendor, and high-throughput crystallography is now widely supported at synchrotrons world-wide. This should therefore become a power tool for fast and robust development of covalent ligands against many different proteins, as demonstrated by the development of two new covalent probes for targets that have lacked inhibitors to date.

## Methods

### Library acquisition and handling

993 compounds were acquired from Enamine (https://www.enamine.net/) as 20 mM DMSO stocks in 96 deep-well plates. A working copy was formatted to 384 well-plates and was kept at room temperature under nitrogen. The rest of the library was aliquoted to 384 well-plates and frozen in −20 °C. Echo 550 liquid handler (Labcyte Inc.) was used to make screening plates with appropriate volumes of compound. Chemical descriptors of the library were calculated using Pipeline-Pilot (Biovia).

### Thiol reactivity assay

50 μM DTNB was incubated with 200 μM TCEP in 20 mM sodium phosphate buffer pH 7.4, 150 mM NaCl, for 5 minutes at room temperature, in order to obtain TNB^2-^. 200 μM compounds were subsequently added to the TNB^2-^, followed by immediate UV absorbance measurement at 412 nm at 37 °C. The absorbances were acquired every 15 minutes for 7 hours. The assay was performed in a 384 well-plate using a Tecan Spark10M plate reader. Background absorbance of compounds was subtracted by measuring the absorbance at 412 nm of each compound in the same conditions without DTNB. Compounds were measured in triplicates.

The data was fitted to a second order reaction equation such that the rate constant k is the slope of ln([A][B_0_]/[B][A_0_]). Where [A_0_] and [B_0_] are the initial concentrations of the compound (200 μM) and TNB^2-^ (100 μM) respectively, and [A] and [B] are the remaining concentrations as a function of time as deduced from the spectrometric measurement. Linear regression using Prism was performed to fit the rate against the first four hours of measurements.

### Electrophile library screen

Plates for electrophile library screens were prepared by combining 0.5 μL of 20 mM stock solution of four or five compounds into one well in a 384 well plate. Incubations were performed at 200 μM for each compound and 2 μM of protein (10 μM for BSA) for 24 hours at 4 °C with moderate shaking. Incubation buffers varied between proteins (PBS pH 7.4 for QSOX1 and BSA; 10 mM HEPES pH 7.5, 0.3 M NaCl, 0.5 mM TCEP for PCAF, UPS8 and NUDT7; 10 mM Tris pH 8.2 500 mM NaCl for PBP3^R504C^; 10 mM Na_2_HPO_4_ pH 7.5 100 mM NaCl 5 mM beta mercaptoethanol for NV3CP; 20 mM Na phosphate pH 7.5 for NNMT and 50 mM NaCl, 20 mM Tris pH 8.0 for K-Ras^G12C^). The reaction was stopped by quenching with formic acid, 0.4% final concentration.

The LC/MS runs were performed on Waters ACUITY UPLC class H, in positive ion mode using electrospray ionization. UPLC separation using C4 column (300 Å, 1.7 μM, 21 mm×100 mm). the column was held at 40 °C, and the autosampler at 10 °C. Mobile solution A was 0.1% formic acid in water and mobile phase B was 0.1% formic acid in acetonitrile. Run flow was 0.4 mL/minutes. Gradient used for BSA was 20% B for 2 minutes increasing linearly to 60% B for 4 minutes holding at 60% B for 2 minutes, changing to 0% B in 0.1 minutes and holding at 0% for 1.9 minutes. Gradient for the other proteins was 20% B for 2 minutes increasing linearly to 60% B for 3 minutes holding at 60% B for 1.5 minutes, changing to 0% B in 0.1 minutes and holding at 0% for 1.4 minutes. The mass data was collected at a range of 750-1550 m/z for NV3CP, K-Ras^G12C^ and NUDT7, 700-1300 m/z for QSOX1, PBP3^R504C^, USP8 and PCAF and 1000-2000 m/z for BSA. Desolvation temperature was 500 °C with flow rate of 1000 L/hour. The voltage used were 0.69 kV for the capillary and 46 V for the cone. Raw data was processed using openLYNX and deconvoluted using MaxEnt.

### Labeling assignment

For each measured well, processed peaks were searched to match the unlabeled protein, common small adducts of the unlabeled protein (that could not be the results of fragment labeling and were seen in the control sample), or labeled protein. Labeling percentage for a compound was determined as the labeling of a specific compound (alone or together with other compounds) divided by the overall detected protein species. Peaks whose mass could not be assigned were discarded from the overall labeling calculation. Wells were flagged if there was no peak of unlabeled protein, undefined peak of over 30% or if there was double labeling of a compound. Flagged wells were manually inspected and the labeling assignment was modified if needed. Wells were regarded as “bad wells” if their LC and MS spectra appeared to be of a degraded protein (low intensity and deformed peak shape) or if after deconvolution there were no clear peaks (high noise levels). All the compounds from bad wells were assigned as no available data in Supp. Dataset 2.

### Quantitative processing of mass spectrometry data

A python script for processing the MaxEnt deconvoluted spectra is supplied as Supp. File 2. Below we briefly outline the logic of the processing.

We first identify the ten highest peaks for each well and discard the rest. The peaks are then normalized from ion counts to percentages, where the highest peak is defined as 100%. The unlabeled protein mass is deduced from a reference well that contains just the protein. Up to four non-compound ‘adducts’ are also assigned from that well (buffer adducts, protein oxidations, etc.) these often keep their proportion when a compound labels the protein.

We discard peaks lower than 10% of the maximum peak. For compound labeling assignment (i.e. not the reference well) we also discard peaks that are less than 100 Da heavier than the unlabeled protein (could not be compound). We iterate the remaining peaks first assigning single modification, prioritizing ‘adducts’. In a second pass we try to assign double labeling (adduct + compound or two compounds). In identifying compound peaks we allowed a ‘noise’ level of ± 4 Da, we note that in rare cases we did identify peaks not corresponding to any compound slightly above this noise level but the automatic processing disregards these. If there are more than 4 peaks exceeding 45%, the well is flagged as a “Bad Well”.

For three of the proteins we identified a second major species: K-Ras^G12C^ ~+48 Da in 25/200 wells, USP8 ~+48 Da in 50/200 wells and NV3CP ~+78 Da in 116/200 wells. In these wells we manually assigned compound identity taking into account both main species of the unmodified protein, as well as compound labeling against other proteins in the benchmark.

### Protein sources, expression and purification

<u>BSA</u> was purchased from MP biomedicals cat. 160069. <u>QSOX1</u> (mouse aa 36-550) was a generous gift from Prof. Deborah Fass (Weizmann Institute) and was produced as described in Grossman *et al.^71^*. <u>NV3CP</u> was produced following the procedure described in Hussey *et al.* ^72^. <u>K-Ras</u>^G12C^ (1-169) was expressed and purified as described in Nnadi *et al.* ^18^.

#### PBP3 ^R504C^

The soluble 50-579 aa fragment lacking the N-terminal transmembrane helix of the *ftsi* gene from *Pseudomonas aeruginosa* PAO1 encoding for the penicillin-binding protein 3 (PaPBP3) was amplified by PCR and subcloned into pET47b using restriction enzymes BamHI and HindIII. The clinical mutation arginine 504 to cysteine was introduced by the Qiaquick protocol. The R504C mutant was expressed and purified as follows: transformed BL21 (DE3) cells were grown in LB media and induced with 1 mM IPTG; protein overexpression was carried out at 18 °C for 16 hours; purification was achieved by reversed Ni2+ affinity chromatography using the N-terminal His6 tag followed by tag cleavage using recombinant HRV 3C protease; the protein was then injected onto a 16/60 HiLoad™ Superdex 200 column (GE Healthcare) and eluted in 20 mM Tris–HCl (pH 8) and 400 mM NaCl.

#### NUDT7

Human NUDT7 (residues 14-235) was cloned into pNIC28-Bsa4 with a TEV-cleavable N-terminal hexahistidine tag. After transformation into *E. coli* (BL21(DE3)-R3), expression was performed in TB auto induction medium (FroMedium), supplemented with 20 g/L glycerol, 50 μg/mL kanamycin and 34 μg/mL chloramphenicol. Cultures were grown for four hours at 37 °C, then the temperature was decreased to 20 °C and the cultures were grown for another 20 hours. Cells were spun at 5,000 rpm for 10 minutes, then resuspended in 0.5 mg/mL lysozyme, 1 μg/mL benzonase, 20 mM imidazole and stirred for two hours at room temperature. 1% Triton X-100 was added and the cells were frozen at −80 °C. On thawing, cells were centrifuged for one hour at 4,000 × g and the supernatant applied to a His GraviTrap column (GE healthcare) equilibrated with binding buffer (10 mM HEPES, 5 % glycerol, 500 mM NaCl, 0.5 mM TCEP, pH 7.5). After washing with binding buffer supplemented to 20 mM imidazole, NUDT7 was eluted with buffer supplemented to 500 mM imidazole. The eluted protein was applied to a PD-10 desalting column (GE Healthcare) and eluted with binding buffer supplemented to 20 mM imidazole. The N-terminal affinity tag was removed by TEV cleavage overnight and uncleaved protein was removed by applying it again to a His GraviTrap column. The flow-through was concentrated and purified further by size exclusion chromatography using a YARRA SEC-2000 PREP column (Phenomenex) equilibrated with binding buffer. Fractions containing protein were pooled, concentrated and stored at −80 °C.

<u>PCAF</u> (aa 23-190) and <u>USP8</u> (aa 705-1081) were produced using the same procedure as NUDT7.

#### NNMT

Wild type hNNMT was cloned into pET-28-TEVH vector harboring an N-terminal His 6-tag. The vector was transformed into E. coli BL21(DE3) cells. Following induction with 1 mM IPTG the culture grew over night at 25 °C. The cells were suspended in lysis buffer (50 mM Tris pH 8, 0.5 M NaCl, 5 mM Imidazole, 2 mM DTT, 5% Glycerol supplemented with protease inhibitor cocktail (Calbiochem), 1 mM PMSF, 0.2 mg/mL lysozyme and 20 μgr/mL DNAse). The cells were lysed using a cooled cell disrupter. The clarified lysate was loaded onto a HisTrap_FF_crude column (GE Healthcare) equilibrated with binding buffer (50 mM Tris pH 8, 0.5M NaCl, 25 mM Imidazole, 5% Glycerol). The enzyme was eluted with the same buffer containing 0.25 M imidazole and injected immediately to a size exclusion column (HiLoad_26/60_Superdex_75). hNNMT eluted in a single peak. The protein was flash frozen using liquid nitrogen in aliquots and kept at −80 °C.

#### OTUB2A

pET20b vector containing Human OTUB2A (residues 1-234) was transformed into the E. coli expression strain BL21(DE3)-R3. Expression was performed in TB medium, supplemented with 0.4% Glucose, 50 μg/mL Ampicillin and 34 μg/mL Chloramphenicol. Cultures were grown at 37 °C till the absorbance at 600 nm of 1.2. The temperature was then decreased to 18 °C and the cultures were grown for further 90 minutes. 10 mM Benzyl alcohol was added and the cells were grown for further 20 minutes before inducing with 0.5 mM IPTG and grown overnight. Following day, cells were centrifuged at 6000 rpm for 30 minutes and the pellet was resuspended in the lysis buffer (50 mM Tris pH 7.5, 500 mM NaCl, 5% Glycerol).

Cells were lysed by sonication for 3 minutes (20 seconds on, 50 seconds off). The lysate was centrifuged at 17000 rpm for 45 minutes. The supernatant was bound to Ni-NTA Agarose beads that were pre-equilibrated with binding buffer (50 mM Tris pH 8, 500 mM NaCl, 20 mM Imidazole), for 1 hour at 4 °C. The beads were washed with binding buffer to pack the column. Protein was eluted with of elution buffer (50 mM Tris pH 8, 500 mM NaCl, 200 mM Imidazole) and supplemented with 0.5 mM DTT. His tag was removed by TEV cleavage while dialyzing OTUB2A overnight in a buffer containing 50 mM Tris pH 8, 500 mM NaCl, 0.5 mM DTT. Dialyzed protein was incubated again with Ni-NTA Agarose beads for an hour. The untagged protein was collected as the flow through, concentrated and purified further by size exclusion chromatography using a HiLoad 26/600 Superdex 75 pg (GE Healthcare Lifesciences) equilibrated with gel filtration buffer (20 mM Tris pH 8, 50 mM NaCl, 5 mM DTT). Fractions containing the protein were pooled, concentrated to 25 mg/mL and stored at −80 °C.

### OTUB2A Crystallization

Microcrystals of OTUB2A were by obtained by mixing 50 nL of OTUB2A (25 mg/mL) with 100 nL of reservoir solution (16% PEG4K, 0.1M HEPES pH 7.0, 8% 2-propanol, 5 mM DTT) in a sitting drop plate at 20 °C. These microcrystals were used for making a seed stock.

In order to first attempt the soaking strategy, 20 nL of the seed stock was used to grow big prism shaped crystals in less than 24 hours. An ECHO 550 acoustic liquid handler (Labcyte) was used to transfer individual fragments from the covalent fragment library to drops containing crystals. Briefly, compound solution was added to each crystallisation drop resulting in a final compound concentration of 4 mM with 20% DMSO, calculated based on the initial drop volume. Crystals were incubated for 2 hours and 24 hours at room temperature. All structures except the compound PCM-0102973 were obtained by co-crystallization.

For co-crystallization of OTUB2A with the compounds of covalent fragment library, 100 μL of the crystallization cocktail (16% PEG4K, 0.1M HEPES pH 7.0, 8% 2-propanol, 5 mM DTT) was dispensed in the reservoir of a sitting drop plate. ECHO 550 acoustic liquid handler was then used to dispense 75 nL of protein and 1-4 mM of the compound on top of the protein drop. The mix was incubated at 20 °C overnight. Next day, 75 nL of the reservoir solution was added on top of the drop along with 20 nL of the seed stock. The plate was incubated at 20 °C and crystals were obtained within 24 hours.

All crystals were harvested with 20% ethylene glycol as cryoprotection and flash cooled in liquid nitrogen. All X-ray diffraction data were collected on the beamline I04-1 at Diamond Light Source (Harwell, UK) unless stated otherwise.

### OTUB2 Structure determination

Diffraction data were automatically processed by software pipelines at the Diamond Light Source ^73^. Initial refinement and map calculation was carried out with DIMPLE ^74^. PanDDA ^75^ was used for hit identification. Further refinement and model building was performed with REFMAC ^76^ and COOT ^77^, respectively. Coordinates and structure factors for all data sets are deposited in the RCSB Protein Data Bank under PDB IDs 5QIO, 5QIP, 5QIQ, 5QIR, 5QIS, 5QIT, 5QIU, 5QIV, 5QIW, 5QIX, 5QIY, 5QIZ. Data collection and refinement statistics are available from the PDB pages.

### OTUB2 inhibition assays

The assays were performed in “non-binding surface flat bottom low flange” black 384-well plates (Corning) at room temperature in a buffer containing 50 mM Tris.HCl, 100 mM NaCl, pH 7.6, 2.0 mM cysteine, 1 mg/mL 3-[(3-cholamidopropyl) dimethylammonio] propanesulfonic acid (CHAPS) and 0.5 mg/mL gamma-globulins from bovine blood (BGG). Each well had a final volume of 20.4 μL. The compounds were dissolved in 10 mM DMSO stocks and appropriate volumes were transferred to the empty plates using a Labcyte Echo acoustic dispenser. A DMSO back-fill was performed to obtain equal volumes of DMSO (400 μL) in each well. 10 mM *N*-ethylmaleimide (NEM) was used a positive control (100% inhibition) and DMSO as negative control (0% inhibition). 10 μL buffer was added and the plate was vigorously shaken for 20 sec. Next, 5 μL OTUB2 (full-length) was added to a final concentration of 25 nM followed by incubation for 30 minutes. or 150 minutes. 5 μL of the substrate (Ub-Rho) was added (final concentration 400 nM) and the increase in fluorescence over time was recorded using a BMG Labtech Clariostar plate reader (excitation 487 nm, emission 535 nm). The initial enzyme velocities were calculated from the slopes, normalized to the positive and negative controls and plotted using GraphPad Prism 7 to obtain the IC_50_ values.

### DUB ABPP assays

All assays were performed in a buffer containing 50 mM Tris, pH 7.4, 5 mM MgCl_2_, 250 mM sucrose, 5 mM DTT, 2 mM ATP. Purified recombinant OTUB2, HEK293T cell lysate (2.5 mg/mL) and HEK293T cell lysate spiked with purified recombinant OTUB2 (0.05 μg/μL and 0.1 μg/μL) were incubated with 50 μM of the inhibitors for 3 hours at 37 °C. Iodoacetamide (10 mM) was used as positive control. Next, Rho-Ub-PRG probe (10 μM) was added and the samples were incubated for another 30 minutes. at 37 °C. Proteins were resolved on a 4-12% NuPage Novex Bis-Tris gel using MOPS running buffer. Rho-Ub-PRG bound DUBs were visualized by fluorescence scanning of the gel on a GE Typhoon GoldSeal FLA9500 scanner (excitation 473 nm) and protein loading was checked by Expedeon InstantBlue staining.

DUB inhibition of NUDT7 inhibitor NUDT7-COV-1 was assessed using a similar method. HEK293T cell lysate was incubated with a concentration series of 0.1-100 μM of the compound and *N*-ethylmaleimide (15 mM) was used as positive control.

### NUDT7 crystallization

NUDT7 crystals were obtained by mixing 100 nL of 30 mg/mL protein in 10 mM Na-HEPES pH 7.5, 500 mM NaCl, 5% glycerol with 50 nL of reservoir solution containing 0.1 M BisTris pH 5.5, 0.1 M ammonium acetate and 6%(w/v) PEG 10,000. Compact, hexagon-shaped crystals with typical dimensions between 50 – 100 μm appeared within several days from sitting drop plates at 20 °C. Co-crystals of NUDT7 in complex with NUDT7-REV-1 and NUDT7-COV-1 were obtained by soaking NUDT7 crystals with a mixture containing 600 nL of 100 mM of the respective compound in DMSO with 1200 nL reservoir solution. Crystals were incubated overnight at room temperature and then harvested (without further cryoprotection) and flash cooled in liquid nitrogen. Crystals of NUDT7 with covalent fragments were grown by mixing 100 nL of 30 mg/mL protein in 10 mM Na-HEPES pH 7.5, 500 mM NaCl, 5% glycerol with 30 nL of 20 mM compound in DMSO in sitting-drop crystallization plates containing 0.1 M BisTris pH 5.5, 0.1 M ammonium acetate and 4 - 16%(w/v) PEG 10,000 in the reservoir at 20 °C. After overnight incubation of protein and compound, 100 nL of reservoir solution and 30 nL of a crystal seed solution obtained from a previous crystallisation experiment, were added to the drop. Hexagon-shaped crystals appeared within several days. Prior to data collection, all crystals were transferred to a solution consisting of the precipitation buffer supplemented with 25% ethylene glycol and subsequently flash cooled in liquid nitrogen. All X-ray diffraction data were collected on beamline I04-1 and beamline I03 at the Diamond Light Source (Harwell, UK).

### NUDT7 structure determination

Diffraction data were automatically processed by software pipelines at the Diamond Light Source ^73^. Initial refinement and map calculation was carried out with DIMPLE ^74^. PanDDA ^75^ was used for hit identification and further refinement and model building was performed with REFMAC 76 and COOT ^77^, respectively. All structure determination steps were performed within the XChemExplorer ^78^ data management and workflow tool.

Coordinates and structure factors for all data sets are deposited in the Protein Data Bank under group deposition ID G_1002045. Data collection and refinement statistics are summarized in Supp. Table 5. The complete PanDDA analysis and all processed data from the NUDT7 fragment campaign (including information about soaked compounds) can be accessed via the ZENODO data repository under DOI <u>10.5281/zenodo.1244111</u>.

### NUDT7 activity assay

Mass spectrometry assays monitoring acetyl-CoA hydrolysis by NUDT7 were performed on a Agilent 6530 RapidFire QTOF Mass Spectrometer in a 384-well plate format using polypropylene plates (Greiner, code 781280) and an assay buffer containing 20 mM HEPES pH 7.5, 200 mM NaCl and 5 mM MgCl_2_. All bulk liquid handling steps were performed using a multidrop combi reagent dispenser (Thermo Scientific, Code 5840300) equipped with a small tube plastic tip dispensing cassette (Thermo Scientific, Code 24073290). For inhibitor IC_50_ determinations an 11-point and 2-fold serial dilution in was prepared from a 50 mM stock solution in DMSO which was transferred to give four replicates using an ECHO 550 acoustic dispenser (Labcyte). The transferred volume was 400 nL giving a final DMSO concentration of 0.4 %. In addition, a DMSO control (400 nL) was transferred into alternate wells in columns 12 and 24 and 50 mM EDTA (NUDT7 inhibitor) was dispensed into alternate wells of column 24 as the background control. 80 μL assay buffer was added to all wells and NUDT7 was prepared to 500 nM (10 X final concentration in assay buffer) and acetyl-CoA was prepared to 200 μM (10 X final concentration in assay buffer). 10 μL NUDT7 was dispensed into half of the assay plate (for two of the compound replicates) and the plate was incubated at room temperature for 30 minutes. 10 μL NUDT7 was then dispensed into the remaining half of the assay plate (for the remaining two compound replicates). 10 μL acetyl-CoA was immediately dispensed into all wells of the assay plate to initiate the reaction and the enzyme reaction was allowed to proceed for 15 minutes. The enzyme reaction was stopped by addition of 10 μL of 50 mM EDTA and the plate was transferred to a RapidFire RF360 high throughput sampling robot. Samples were aspirated under vacuum and loaded onto a C4 solid phase extraction (SPE) cartridge equilibrated and washed for 5.5 sec with 1 mM octylammonium acetate in LCMS grade water to remove non-volatile buffer components. After the aqueous wash, analytes of interest were eluted from the C4 SPE onto an Agilent 6530 accurate mass Q-TOF in an organic elution step (85% acetonitrile in LC-MS grade water). Ion data for the acetyl-CoA and hydrolysed product were extracted and peak area data integrated using RapidFire integrator software (Agilent). % conversion of substrate to product was calculated in excel and IC_50_ curves generated using Graphpad prism version 7.0. The assay had a Z score of 0.79 with the 30 minutes pre-incubation and 0.75 without pre-incubation.

### NUDT7 Thermal shift assay

5 μM NUDT7 was incubated prior to the measurements with 200 μM of compound in 10 mM HEPES pH 7.5, 0.3 M NaCl, 0.5 mM TCEP for 24 hours at 4 °C. 1 μL of 5x SYPRO Orange (sigma) was added to 19 μM of incubated protein in MicroAmp Fast Optical 96-Well Reaction Plate sealed with MicroAmp Optical Adhesive Film. Measurements were performed using StepOnePlus rtPCR from 25 °C to 95 °C with 0.3 °C steps. T_m_ was determined using the StepOne Software v2.3. Reported T_m_ was calculated as the average of three triplicates for each compound.

### NUDT7 cellular thermal shift assay

HEK293 cells were cultured at 37 °C in a humidified 5% CO_2_ atmosphere in DMEM supplemented with GlutaMAX and 10% FBS. Cells were grown in T175 flasks until around 70% confluent and transfected with Flag-NUDT7 using Lipofectamine2000 transfection reagent. Twenty-four hours post-transfection, cells were detached and 13 × 10^6^ cells were seeded in T75 flasks for treatment and control sample, respectively. After 24 hours cells were treated either with DMSO or 20 μM NUDT7-COV-1 for 30 minutes at 37 °C. Cells were harvested, washed with PBS and aliquoted into PCR tubes. PBS was removed by centrifugation (300 × g, 3 minutes, RT). Cell pellets were heated to temperatures ranging from 37 to 73 °C with 4 °C increments for 3 minutes (UNO96, VWR), cooled down to room temperature for 3 minutes and then transferred onto ice. Lysis was performed in lysis buffer (50 mM Tris pH 7.5, 0.8% v/v NP-40, 5% v/v glycerol, 1.5 mM MgCl_2_, 100 mM NaCl, 25 mM NaF, 1mM Na_3_VO_4_, 1 mM PMSF, 1 mM DTT, 10 μg/mL TLCK, 1 μg/mL leupeptin, 1 μg/mL aprotinin, 1 μg/mL soy bean trypsin) by three freeze-thaw cycles in liquid nitrogen. The resulting lysates were centrifuged at 21,000 g for 20 minutes at 4 °C to remove aggregated proteins. Protein concentration was determined for the soluble fraction followed by by SDS-PAGE analysis followed and Western blotting. After transfer the nitrocellulose membrane was blocked with blocking buffer (5% (m/v) BLOT-QuickBlocker (Merck) in PBST (Phosphate-buffered saline with 0.05% (v/v) Tween 20)) and probed with primary antibody (mouse anti-FLAG (Merck, F3165) 1:1,500 in blocking buffer) overnight at 4 °C and secondary antibody (goat anti-mouse Alexa Fluor 750 (Life Technologies, A-21037) 1:10,000 in blocking buffer) for 1 hour at RT. Blots were imaged on an Odyssey CLx imager (LI-COR).

### Cell viability assay

HEK293, HB2 or CCD841 cells grew in either RPMI or DMEM mediums supplemented with 10% FCS, 1% PS and 1% L-Glutamine (all from Biological Industries). Exclusion of Mycoplasma contamination was monitored and conducted by test with Mycoalert kit (LONZA). Cells were trypsnized, counted, and 1000 cells/well were plated in 50μL of growth medium into 384-well white TC plates (Greiner) using MultiDrop 384 (Thermo Scientific) Washer Dispenser II. Number of viable cells was monitored using CellTiter-Glo^®^ Luminescent kit (Promega) in accordance to the manufacture protocol. Luminescence was measured using luminescence module of PheraStar FS plate reader (BMG Labtech). Data analysis was performed using GeneData 12 analytic software.

### NNMT activity assay

Compounds transferred into black microplates (Greiner 784900) using Labcyte Echo acoustic dispensing. Assay ready plates were then sealed with heat seals. If not used immediately, plates were frozen at −20 °C and held in polypropylene boxes with silica-gel desiccant.

Reagents were obtained as follows: Nicotinamide (Sigma 47865-U), SAM (Sigma A7007), SAH-FITC (Axis Shield, RPBB350), Anti-SAH (Axis Shield, RPBB278). All liquid handling was done with a GNF washer/dispenser II. All reagents were prepared in 20 mM phosphate buffer, pH=7.6. 6 μL of 3X NNMT (120 nM) was added to assay plates and incubated for 10 minutes at room temperature. 6 μL of 3X substrate mixture (120 mM Nicotinamide, 6 μM SAM) was added and incubated for 1 hour at 30 °C. 6 μL of 3X detection mixture (150 ng/mL SAH-FITC and 30 μg/mL anti-SAH antibody) was prepared and incubated for 30 minutes at room temperature before adding to reactions. Plates were further incubated for another 2 hours at 30 °C, protected from light. Fluorescence polarization reaction were read in BMG Pherastar FS using a 485/520/520 nm module. Data was normalized to DMSO (100%) and no enzyme (0%) controls using Genedata Screener software.

## Acknowledgments

N.L. is the incumbent of the Alan and Laraine Fischer Career Development Chair; N.L. would like to acknowledge funding from the Israel Science Foundation (grant No. 1097/16), The Rising Tide Foundation, The Israel Cancer Research Foundation and the Israeli Ministry of Science and Technology (grant No. 3-14763). This work was supported by the Dutch Organization for Scientific Research NWO (VICI grant 724.013.002 to H.O.) We thank Yves Leestemaker and Jin Gan for assistance with the DUB ABPP assays. The SGC is a registered charity (number 1097737) that receives funds from AbbVie, Bayer Pharma AG, Boehringer Ingelheim, Canada Foundation for Innovation, Eshelman Institute 22 for Innovation, Genome Canada through Ontario Genomics Institute [OGI-055], Innovative Medicines Initiative (EU/EFPIA) [ULTRA-DD grant no. 115766], Janssen, Merck KGaA (Darmstadt, Germany), MSD, Novartis Pharma AG, Ontario Ministry of Research, Innovation and Science (MRIS), Pfizer, So Paulo Research Foundation-FAPESP, Takeda, and Wellcome [106169/Z/14/Z]. We thank Prof. Deborah Fass for the generous gift of QSOX1 protein.

